# Highly Localized Responses to Salinity and Drought in Rice Roots

**DOI:** 10.1101/2023.10.29.564585

**Authors:** Rukaya Amin Chowdery, H E Shashidhar, M K Mathew

## Abstract

Drought and salt stress are first sensed by the root system of plants. Many physiological responses, including variation in stomatal conductance, are regulated by the stress hormone ABA which is generated in the root and sent up to the shoot which then synthesizes additional ABA to sustain the response. To address whether responses are systemic we have used a split root system, wherein roots are divided into two and each half treated independently to water, salt or drought. Four varieties were examined – the salt tolerant Pokkali, the drought tolerant ARB6 and two sensitive varieties Jaya and IR-20. ABA concentrations in the xylem sap increased dramatically after Day 1 in all four cultivars in response to stress on at least one side the of the split root system. The sensitive varieties appeared to derive much of their nutrition and fluid from the watered side when subjected to asymmetric conditions, whereas roots on the stressed side of tolerant varieties underwent anatomical and physiological modifications facilitating fluid uptake and maintenance of xylem sap flow under these conditions.

## Introduction

Salinity and drought are highly deleterious stresses that severely impair plant growth and crop production (Wang et al., 2003; Tuteja, 2007; Qin et al., 2011). Plant roots are the first to perceive and integrate changes in soil (Malamy, 2005; Sánchez-Calderón et al., 2013). Modulation of the architecture of the root system has been reported to facilitate location of water and nutrients in water deficit conditions (Lynch, 1995). Remodeling of internal barriers affects root hydraulics and solute loading into the xylem stream. We have asked whether roots adopt global or local strategies to cope with stress conditions in the soil and used the split root system to address this question. Split root and grafting experiments have been instrumental in dissecting the effects of environmental and endogenous factors that affect responses to abiotic stresses.

The split root system has been used to demonstrate that roots extract water from both macropores and micropores alike (Emerman, 1995; Emerman and Dawson, 1997). When wheat, maize and Arabidopsis were subjected to an uneven distribution of nitrogen, the length, number of lateral roots and metabolic activity in the nitrogen supplied side increased (Hackett, 1972; Granato and Raper, 1989; Zhang and Forde, 1998; Zhang and Forde, 2000; Wang et al., 2002). Other studies show that non-uniform salinity in soil increased the nutrient uptake on low salinity side of the root in cucumber (Sonneveld and De Kreij, 1999), more yield of tomato in plants where half of the root received water treatment compared to when both sides of the root were treated with salinity (Tabatabaie et al., 2003) and high plant biomass and yield in cotton where both sides of the root were treated with low salinity condition (Dong et al., 2010). Besides maintaining water uptake during water deficit conditions by modulating root architecture, plant roots also have the ability to communicate the information to the shoot (Gollan et al., 1986; Passioura, 1988).

Long distance root-to-shoot signaling mechanisms are important in plants to survive under unfavorable conditions. In response to different environmental cues like drought, salt and cold, abscisic acid (ABA), a well characterized phytohormone which is involved in root to shoot signaling (Zhang and Davies, 1987; Dodd, 2005) has been found to increase significantly (Shinozaki and Yamaguchi-Shinozaki, 2000). The main function of ABA is to regulate plant water balance and osmotic stress tolerance (Zhu, 2002; Hirayama and Shinozaki, 2007; Wasilewska et al., 2008; Santner et al., 2009; Cutler et al., 2010) by inducing stomatal closure which in turn helps adapting plants towards severe water shortage (Davies and Zhang, 1991; Finkelstein et al., 2002; Hirayama and Shinozaki, 2007; Nambara and Marion-Poll, 2010). In heterogeneous soil moisture condition, leaf ABA concentration was reported to be high in barley plants with more roots in drying side of the chamber (Martin-Vertedor and Dodd, 2011), these findings also prompted the question whether or not the root-to-shoot signaling in rice would be dependent on equal root distribution when faced with uniform and non-uniform drought and salinity stress in split root setup. Also to check whether equal distribution of roots on both sides of the split root setup would result in increased ABA concentration in leaves when subjected to these conditions in rice.

Various grafting and split root experiments have been conducted in many plants like maize (Dodd et al., 2008), tomato (Holbrook et al., 2002) and barley (Dodd et al., 2002; Martin-Vertedor and Dodd, 2011) to understand the response of plants under heterogeneous soil moisture conditions, but no studies have been reported in rice. In the present study, four rice cultivars were selected for comparison based on their sensitivity to drought and salinity. A split-root system was established to study morphological and physiological responses of these rice cultivars to both uniform well-watered (W/W), drought (D/D), salinity (S/S) and non-uniform W/D, W/S and D/S conditions of drought and salinity stresses, focusing on: (i) root architecture and anatomy on both sides of the split root; (ii) water uptake by cultivars under uniform and non-uniform stress; (iii) suberization patterns in roots under these conditions; (iv) effect of these stresses on canopy temperature of these cultivars; (v) effect on xylem sap and leaf ABA concentration; and (vi) the effect of ABA on stomatal conductance.

## Materials and Methods

### Split-root experiments

For this experiment, two PVC pipes of diameter 16cm and length 80cm were joined together using tape to create two chambers and then filled with a mixture of soil and manure in the ratio 2:1 respectively and pH 5.7. To achieve the field like compaction of soil in PVC pipes the method described by (Shashidhar et al., 2012) was used. Seeds of four cultivars of rice Pokkali (salt tolerant), ARB6 (drought tolerant), IR-20 (salt sensitive) and Jaya (moderately salt sensitive) were used. One month old plants of these cultivars which were grown in well-watered condition were transferred to split-root experimental setup (Fig. 1a). The roots of these plants were carefully divided into two equal halves and planted into split-root pipes (Fig. 1b). For these plants to become healthy and stable, they were further grown and watered regularly for 33 days. The PVC pipes were arranged in a randomized complete block design in field and divided into 6 batches, one batch for each condition. These plants which were about 63 days old were then subjected to 6 different conditions. From 64^th^ day onwards, one of the batches was subjected to water/water (W/W) condition, and other five batches were subjected to drought/drought (D/D), salt/salt (S/S), water/drought (W/D), water/salt (W/S) and drought/salt (D/S) conditions respectively for a week. In total 864 pipes were used with 144 pipes in each batch and 36 replications per variety using 6 replicates for each analysis. The side of the pipe marked water was watered regularly whereas the drought side of the pipe was not irrigated untill the soil reached about 30-40% of field capacity and salinity side was exposed to 150mM NaCl stress for one week. By the end of the experiment the plants were 70 days old. Soil moisture content was monitored using moisture meter (Procheck, Decagon Devices Inc., USA) (Fig. 1c), where the probe was directly inserted into the soil from top of the pipe. Electrical conductivity of soil in PVC pipes after irrigation with normal and saline water of 150mM NaCl concentration was also measured (Fig. 1d).

**Figure 1.**
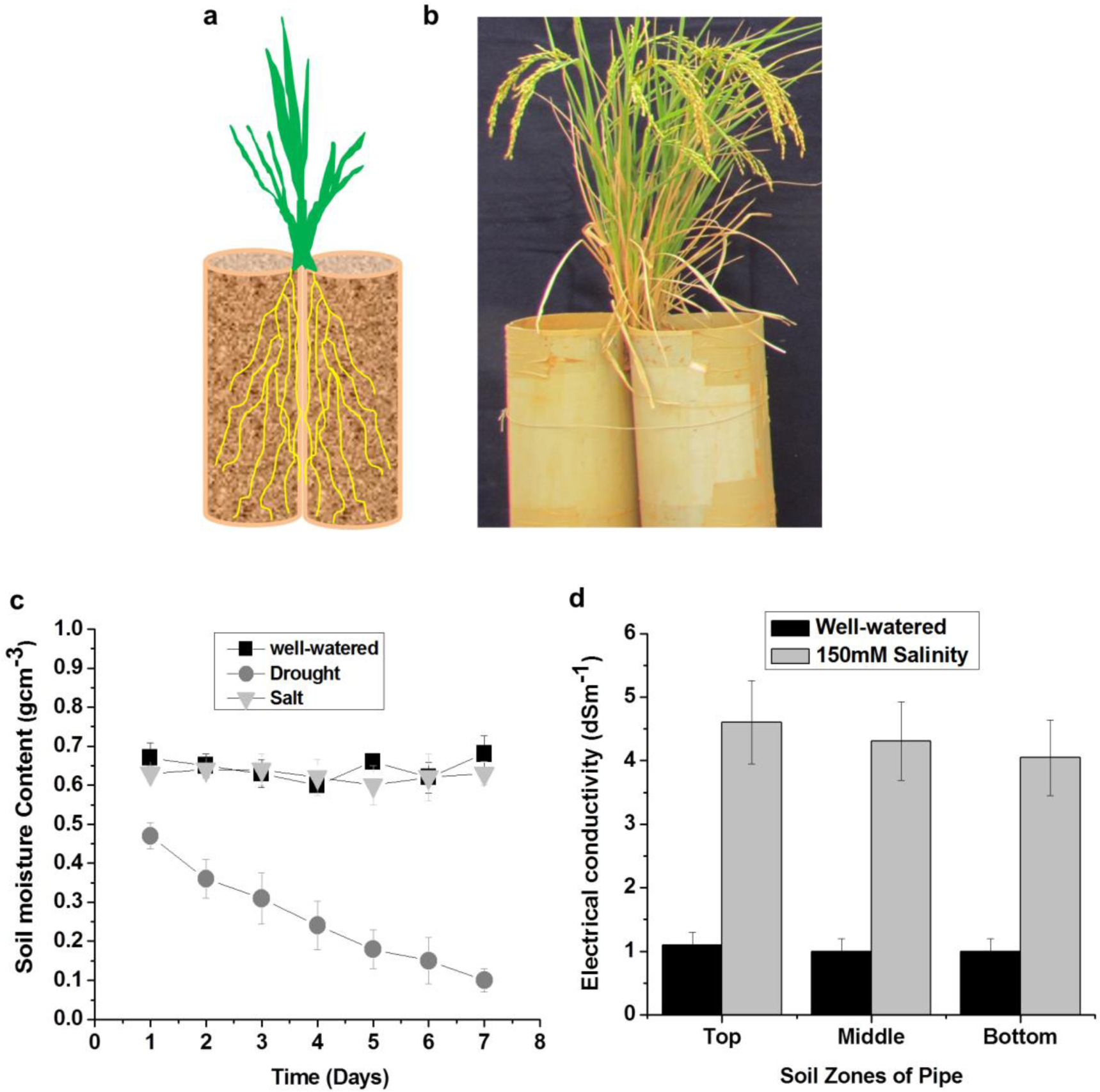
(a) Schematic split-root experiment showing equal halves of root growing in separate chambers. (b) Plants grown in PVC pipes in split-root experimental condition. (c) Soil moisture was measured directly inserting a probe into soil under control (well-watered), drought and Salt conditions. (d) Electrical conductivity of soil in PVC pipe at top, middle and bottom zones under well-watered and 150mM salinity condition.

### Analysis of plant morphology and biomass

After one week of stress, the soil column along with the whole plant was taken out of the pipe. Roots were carefully washed free of soil. The total shoot and root lengths were measured and number of tillers and lateral roots was also counted manually. At this stage the fresh weight of the plants was recorded and dry weight after 48 hours of oven-drying at 70°C.

### Estimation of crop canopy air temperature difference (CCATD)

After one week of stress, ambient and canopy temperature of plants in all six conditions was recorded by using AGRI-THERM III^TM^ infra-red sensing instrument (Everest Interscience Inc., USA) at mid-day on the 70^th^ day. The difference in the two readings gives the Crop Canopy Air Temperature Difference (CCATD).

### Measurement of xylem sap exudation

Sap measurements were recorded using the method described (Morita and Abe, 2002; Henry *et al*., 2012) with some modifications. Rice varieties were grown in six replicates in split-root PVC pipes and exposed to six different conditions as described above. At 06:00 pm on the 70^th^ day, shoots were cut ∼5 cm above the soil surface. Cut stems attached to the root system were covered with pre-weighed blotting paper, then covered with polyethylene wrapper, which was in turn tightly sealed at the base with a rubber band (to avoid evaporation of the sap). The setup was left for 12 hours, following which blotting papers were removed and immediately weighed to quantify the amount of xylem sap sent up by the root system. The values were normalized to the root mass of the plant from which sap was collected.

### Root microscopy

Roots from both sides of W/D and W/S condition were washed carefully. Thin sections (200µm) from different zones of the root like Base (25 cm), Middle (15 cm) and Tip (5 cm) were cut using a Mcllwain tissue chopper (The Mickle Laboratory Engineering Co. Ltd, UK). Sections were stained as described by (Brundrett et al., 1988) with 0.1% (w/v) berberin hemisulphate for one hour followed by 0.5% (w/v) aniline blue for 30 minutes. Stained sections were imaged under Olympus FV1000 confocal microscope using 488nm excitation and 510-540nm emission. For counting and estimation of number of cells with or without suberin deposits, stained sections were observed using UV light under Nikon fluorescent microscope. Suberin was seen as a fluorescent light yellow band shaped structure deposited around the endodermal and exodermal cells of the root, whereas Passage cells (PCs) showed no tangential suberin deposition.

### Measurements of ABA in leaves and xylem sap

To measure changes in ABA, about 3 leaves were collected from 3 plants of each cultivar on Day 0 (one day before subjecting the plants to 6 different conditions), Day 1, Day 2, Day 3, Day 4 and Day 7. The xylem sap was collected using the blotting paper method as described above. Leaves and blotting papers were wrapped in aluminum foil and labeled, then immediately frozen in liquid nitrogen and brought to lab. Leaf tissue and blotting papers were thawed and ABA was extracted with hot water as described by (Loveys and Van Dijk, 1988). ABA was assayed in triplicates by an indirect ELISA method as described by (Walker-Simmons, 1987) using a monoclonal antibody (Idetek).

### Measurement of stomatal Conductance

Stomatal conductance was measured using a porometer (Delta-T) on Day 0, Day 1, Day 2, Day 3, Day 4 and Day 7 using 3 plants per cultivar between 12.00 h to 14.00 h on 6 leaves per plant.

### Statistical analysis

The data in the figures have been presented as the mean values ± SE; n = 6, whereas the ABA data is presented as the mean values of ± SE; n = 3. Student’s t-test was used to estimate the significant differences at P < 0.01 and P < 0.05 between six conditions.

## Results

### Root analysis

After exposing the plants to uniform and non-uniform water, drought and salinity stress for one week, where soil moisture of well-watered and salinity side of the pipe was maintained almost at same levels but higher than drought side where it was very low and decreasing with each day of stress (Fig. 1a, b, c). The electrical conductivity of the soil column in the pipe exposed to 150mM salt stress was about 4 dS from top to bottom (Fig. 1d), which is considered as a critical salt stress level for glycophytes such as rice. It was quite intriguing to observe that all four cultivars showed symmetric growth of roots in both sides of the PVC pipe when exposed to uniform stress (Fig. 2a, b, c) as opposed to non-uniform condition (Fig. 2d, e, f) where the growth was asymmetric. After measuring the root lengths from both sides of the pipe in all six conditions it was observed that the roots of ARB6 were longer in W/W condition followed by Pokkali, Jaya and IR-20 (Fig. 3a). Interestingly, the roots of both tolerant Pokkali and ARB6 growing in the drought side of the pipes in D/D and W/D condition were found to be longer than the roots in water or salt side of the pipe (Fig. 3a). The more prominent increase of the root length in drought condition was observed in ARB6. In case of S/S condition, root lengths of Pokkali in both sides of the pipes was almost same but more than W/W condition, whereas in W/S and D/S condition the root length was observed to be more in salt side of the pipe than the other side (Fig. 3a), which was not observed in case of ARB6 where the root length was observed to be the same on both sides of the pipe in both these conditions, but more under W/S than D/S condition (Fig. 3a). Jaya showed no difference in root lengths in all six conditions except for the S/S where roots showed asymmetric growth and D/S condition where the root on salt side of the pipe had not grown longer as that of the root in drought side of the pipe (Fig. 3a). IR-20 showed no difference in root length in W/W, D/D and S/S condition, though the root lengths in D/D and S/S condition was same but slightly less than W/W condition (Fig. 3a). It was also observed to have grown longer roots on water side of the pipes in W/D and W/S condition than the other side of the pipe. The root length of IR-20 in D/S condition was same on both sides of the pipe and was very less compared to other five conditions (Fig. 3a).

**Figure 2.**
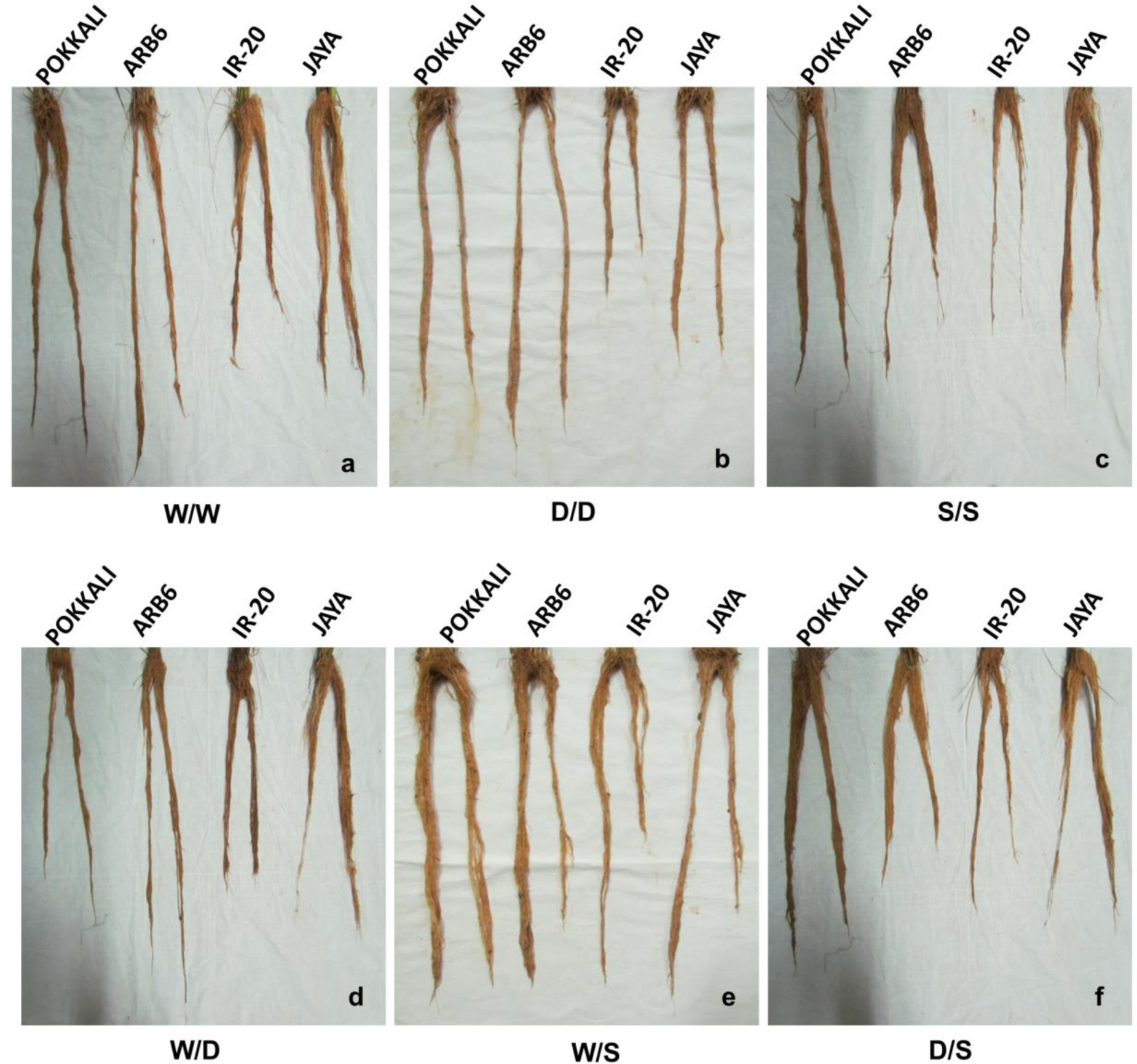
Roots of four rice varieties after the experiment. Plants were grown in PVC pipes for 63 days in split root pipes and were subjected to six different conditions (a) water/water (W/W), (b) drought/drought **(0/0),** (c) salt/salt (S/S), (d) water/drought (W/0), (e) water/salt (W/S) and (f) drought/salt (0/S) for a week.

**Figure 3.**
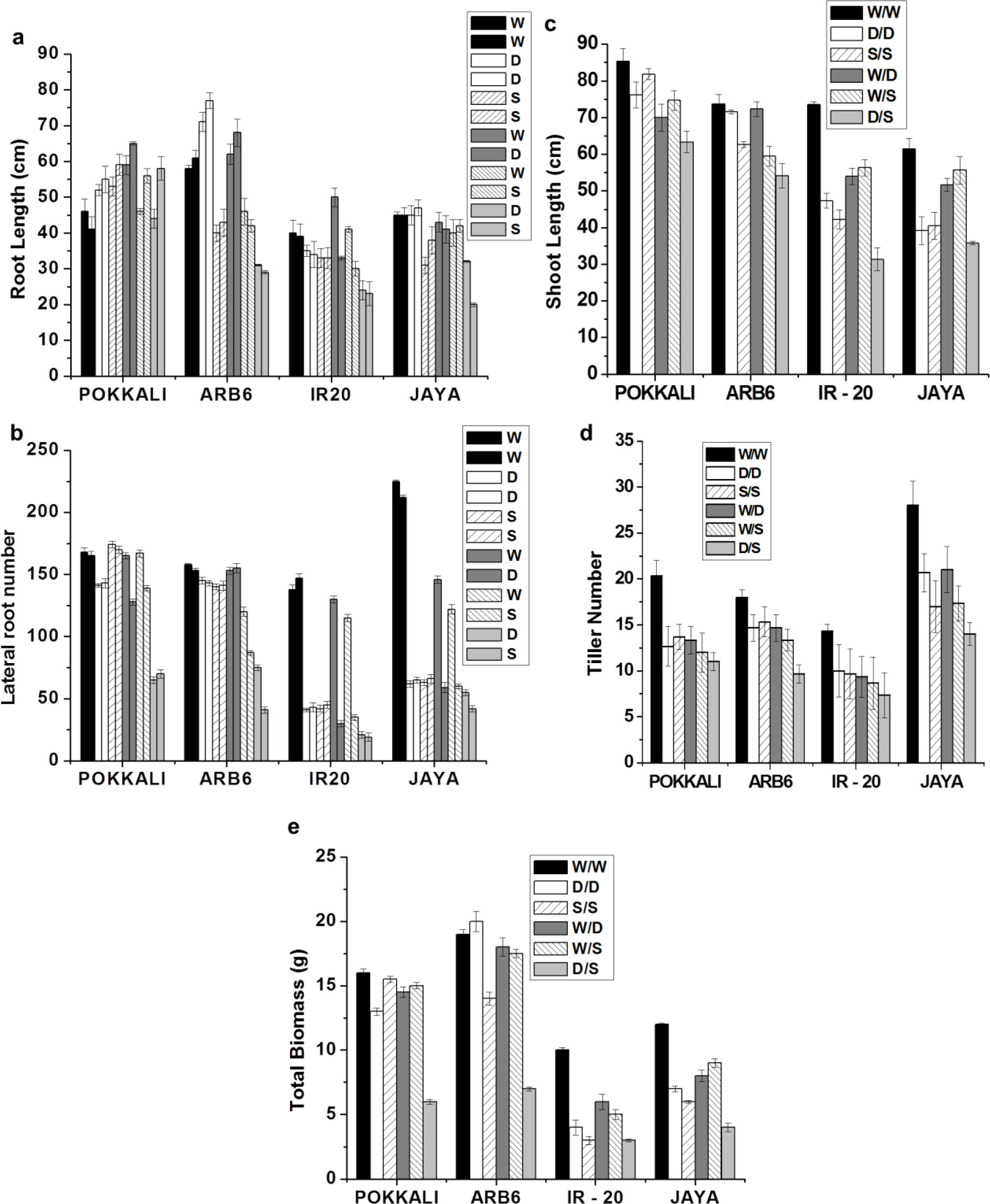
Plants were grown in PVC pipes for 63 days and subjected to six different conditions water/water (W/W), drought/drought (D/D), salt/salt (S/S), water/drought (W/D), water/salt (W/S) and drought/salt (D/S) for a week and following measurements were taken (a) root length in both sides of the pipe in all six stress conditions (b) lateral root number in both sides of the pipe in all six stress conditions (c) shoot length (d) Tiller number (e) total biomass. Data represents mean ±SE, n = 6.

Under W/W condition, Jaya was observed to possess highest number of lateral roots in both sides followed by ARB6, Pokkali and IR-20 the least (Fig. 3b). Interestingly the number of lateral roots possessed by all four varieties changed with different conditions they were subjected to. Under D/D condition, the number of lateral roots in both Pokkali and ARB6 were less than W/W condition on both sides, with more number observed in case of ARB6 than Pokkali whereas, number of lateral roots found in either Jaya or IR-20 was very less (Fig. 3b). Under S/S condition, little increase in lateral root number was observed only in Pokkali than the other three varieties (Fig. 3b). Under W/D condition, the lateral root number in ARB6 was almost similar to that in W/W condition on both sides. Surprisingly, the number of lateral roots possessed by Pokkali, Jaya and IR-20 were more on water side than the drought side of the pipe in W/D condition (Fig. 3b). Under W/S condition, all four varieties possessed more roots on water side than the salt side of the pipe with highest root number possessed by Pokkali than the other three varieties (Fig. 3b). The number of roots in all four varieties under D/S condition were less compared to W/W condition (Fig. 3b).

### Shoot analysis

Longest shoot was observed in Pokkali, ARB6, IR-20 and Jaya in the order under W/W condition (Fig. 3c). The shoot length was observed to decline in all four varieties under other five conditions where marginal decline was observed in case of Pokkali and ARB6 except under D/S condition (Fig. 3c). In Jaya, the decrease in shoot length was more under D/D, S/S and D/S condition than the W/D and W/S condition, whereas the decline was more dramatic in case of IR-20 under all five conditions compared to W/W condition (Fig. 3c).

Under W/W condition, highest number of tillers was found in Jaya, Pokkali, ARB6 and IR-20 in the order (Fig. 3d). The tiller number was found to decrease in all four varieties after they were subjected to D/D, S/S, W/D, W/S and D/S conditions (Fig. 3d).

### Total Biomass

Under W/W condition, ARB6 attained highest total biomass followed by Pokkali, Jaya and IR-20 in decreasing order (Fig. 3e). Only in case of ARB6, the total biomass was observed to increase under D/D condition compared to other three varieties where it was found to decrease (Fig. 3e). Under S/S condition, there was no difference in the total biomass of Pokkali compared to W/W condition but the biomass of other three varieties under S/S condition had decreased (Fig. 3e). Pokkali and ARB6 were found to maintain the biomass same under W/D and W/S as that of W/W condition, whereas, Jaya and IR-20 showed reduction in biomass under these conditions (Fig. 3e). The total biomass had dramatically reduced in all four varieties with ARB6 and Pokkali still managing to maintain high biomass than Jaya and IR-20 under D/S condition (Fig. 3e).

The overall root and shoot morphological analysis indicates that the tolerant varieties such as Pokkali and ARB6 performed well in terms of maintaining the healthy morphology, growth of roots, shoots and biomass under W/W, D/D, S/S, W/D and W/S conditions where they were subjected to either well-watered or drought or salinity or the mixture of these conditions. But when subjected to D/S condition where both drought and salinity stress was given, they exhibited poor morphology. Both Jaya and IR-20 did well under W/W condition, while, Jaya showed moderate morphological performance under W/D and W/S conditions, whereas, under D/D, S/S and D/S conditions both Jaya and IR-20 displayed poor morphology.

### Canopy temperature

Rice plants are in continuous need of water for growth and other metabolic activities. During water limiting conditions such as drought and salinity, they should be able to meet the need either by reducing the rate of transpiration or accessing the additional sources of water by growing dense and longer roots. The fact that the canopy temperature of plants that are actively transpiring would be significantly lower than the ambient, therefore, recording canopy temperature of plants subjected to W/W, D/D, S/S, W/D, W/S and D/S appeared to be a surrogate measure to understand their water status. The canopy temperature of ARB6 was about 7 °C below ambient under W/W condition and that of Pokkali, Jaya and IR-20 was observed to be about 6 °C, 4 °C and 2.5 °C respectively (Fig. 4a), suggesting adequate access of these plants to water. While Pokkali and ARB6 exhibited modest increase in canopy temperatures under D/D, S/S, W/D and W/S conditions, Jaya displayed dramatic increase under D/D and S/S condition, with slight increase under W/D and W/S condition (Fig. 4a). On the other hand, the canopy temperature of IR-20 under D/D and S/S condition was about 1 °C and 0.5 °C more than ambient, whereas under W/D and W/S condition IR-20 exhibited the canopy temperature more or less equal to that of ambient (Fig. 4a). Under D/S condition, the canopy temperature of ARB6 and Pokkali had significantly increased, while in case of both Jaya and IR-20, it had increased about 1°C more than ambient (Fig. 4a).

**Figure 4.**
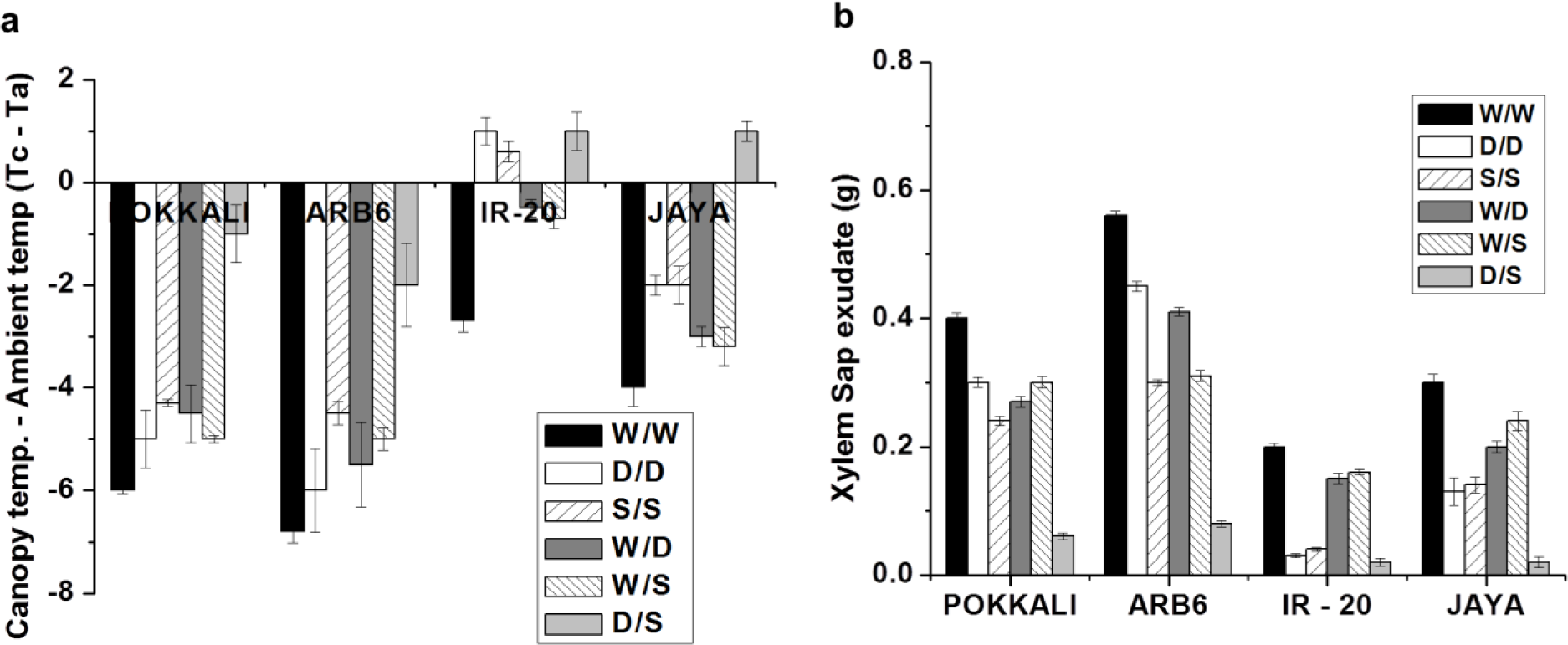
Plants were grown in PVC pipes for 63 days and subjected to six different conditions for a week and following measurements were taken (a) Crop canopy air temperature difference (CCATD) in four rice varieties. Ambient and canopy temperature was measured by using infra­ red sensing instrument. Temperature of canopy was estimated by taking the difference of canopy and ambient temperature (Tc - Ta). (b) Xylem sap exudation. Stems of control and drought plants devoid of shoot were covered with pre-weighed blotting paper for 12 hours and paper was weighed again to estimate the amount of xylem sap absorbed by blotting paper. Data represents mean± SE, n = 6.

### Xylem sap exudation

In order to estimate root-driven xylem flow that would be required to support the observed canopy temperature differences, we measured the amount of xylem sap exuded over a 12 hour period. The root driven xylem sap exudation rates under W/W condition in ARB6 were higher than Pokkali, Jaya and IR-20 in the order (Fig. 4b). Though the amount of xylem sap exudated under D/D, S/S, W/D, W/S and D/S conditions reduced in all four varieties and followed the same trend as that in W/W condition were exudation rate was higher in ARB6 than Pokkali, Jaya and IR-20 (Fig. 4b). In case of ARB6 and Pokkali, the root-driven xylem sap exudation had declined to less than half under D/S condition, whereas, the xylem sap exudation had essentially stopped in IR-20 and Jaya compared to W/W condition, (Fig. 4b).

### Suberin deposition under W/D condition

Pattern of suberin – a hydrophobic barrier deposition in endodermis and exodermis plays an important role in maintaining root hydraulics under drought and salinity stress and therefore, is critical for delivering fluid to the growing shoot. Pokkali – a salt tolerant variety has been shown to deposit extensive suberin barriers under salinity stress (Krishnamurthy et al., 2009; 2011). These results prompted to examine the hydrophobic barriers on the endodermis and exodermis (Fig. 5 and Fig. 6) in all four varieties under W/D and W/S condition to check whether or not these varieties deploy different strategies of suberin deposition in roots on both sides of the pipe.

**Figure 5.**
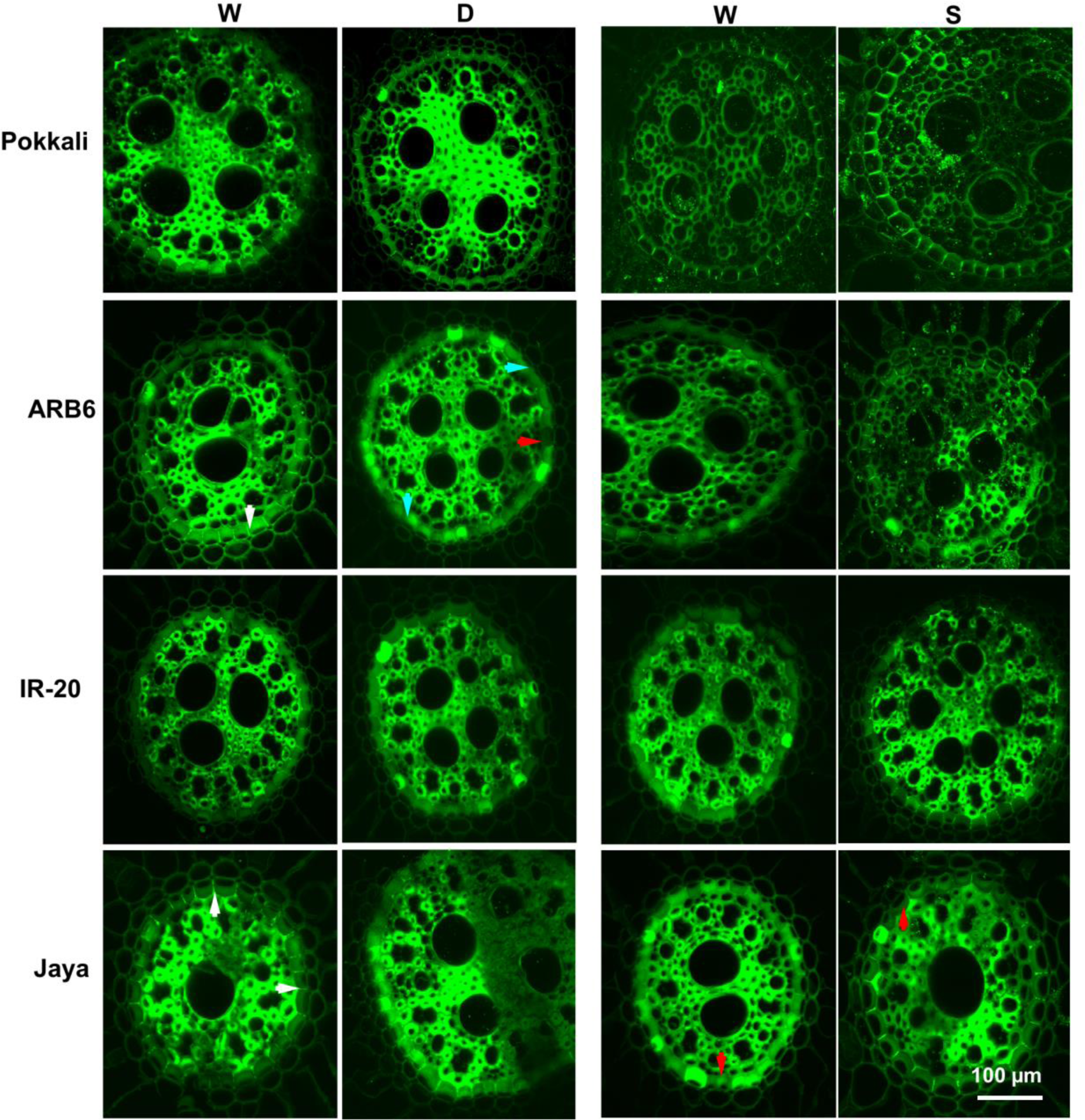
Images of root sections showing unsuberized cells, passage cells and suberin deposits in root endodermis of plants grown in PVC pipes for 63 days and subjected to water/drought (W/D) and water/salt (W/S) condition for a week. Roots were washed, divided into three zones (Tip - 5cm, Mid - 15cm and Base - 25cm) from the root tip and cut into thin sections (200µm), then stained with berberine-aniline blue and imaged using Olympus FV1000 confocal microscope, 488nm laser was used for excitation. Red, white and cyan arrows show passage cells (cells with no tangential suberin deposition), suberin deposits and unsuberized cells respectively. Scale bar 100µm

**Figure 6.**
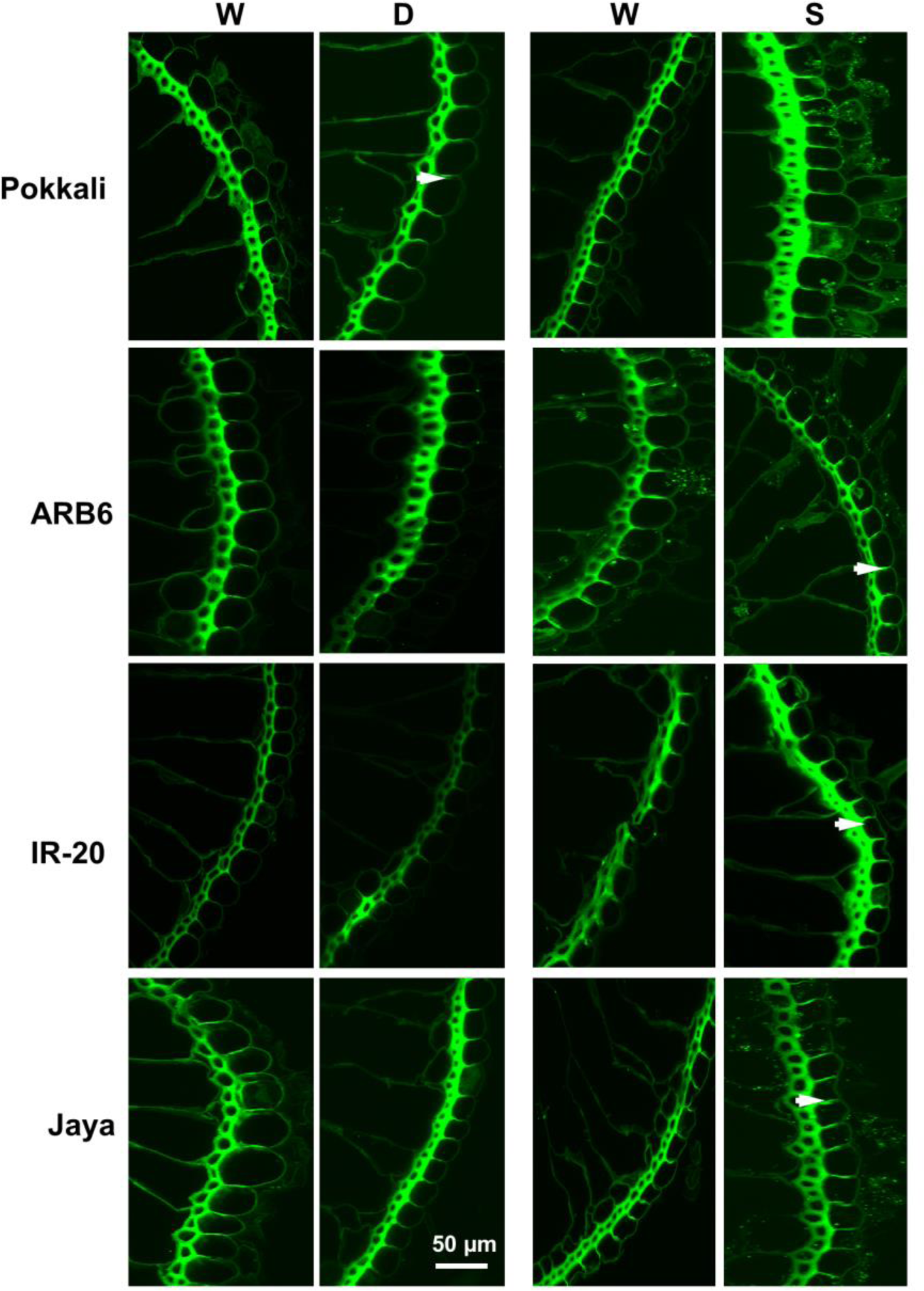
Images of root sections showing unsuberized cells, passage cells and suberin deposits in root exododermis of plants grown in PVC pipes for 63 days and subjected to water/drought (W/O) and water/salt (W/S) condition for a week. Roots were sectioned, stained and imaged as in Figure 9. White arrows show suberin deposits. Scale bars 50µm

Following W/D condition, the percentage of suberized cells in both water and drought side of all four varieties was high at base region of the roots than mid and tip region in decreasing order (Fig. 7a, d, g). The percentage of suberized cells in water side of the Pokkali and ARB6 root endodermis was comparable at the base region but greater than Jaya and IR-20, whereas at mid and tip region it was higher in Pokkali than the other three varieties (Fig. 7a, d, g). Interestingly, the percentage of suberized cells in endodermis on drought side of the root decreased in all four varieties and the decline was more in ARB6 at base, mid and tip region than Pokkali, Jaya and IR-20 (Fig. 7a, d, g).

**Figure 7.**
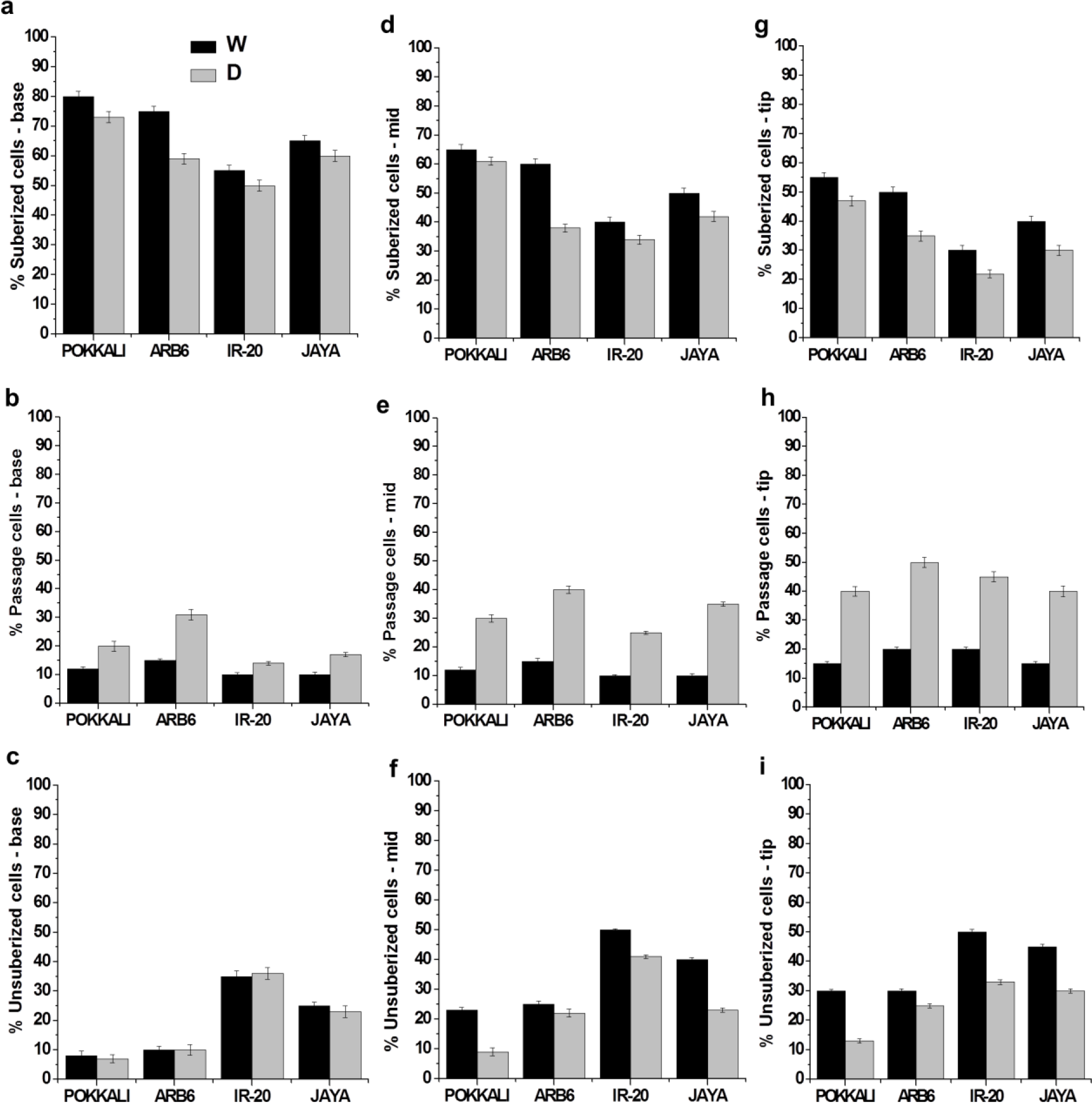
Quantification of suberization patterns under water/drought (W/D) condition at (a - c) base, (d - f) mid and (g - i) tip zone of the root endodermis to show suberized cells (having suberization on both sides and tangential areas), passage cells (with no tangential suberization) and unsuberized cells (without any suberin deposition). Data represents mean (±SE; n = 6)

The percentage of passage cells increased from base to tip of the root in all varieties and was high in drought side of the root than the water side (Fig. 7b, e, h). The highest percentage of passage cells was possessed by tolerant ARB6 than Pokkali, Jaya and IR-20, which was interestingly true at the base region of ARB6 also (Fig. 7b, e, h). Passage cells have the potential of symplastic uptake of fluid, increasing their number by tolerant varieties on drought side of the root suggests their possible assistance in more water uptake.

The percentage of unsuberized cells at base, mid and tip region in the Pokkali and ARB6 endodermis was comparable but less than IR-20 and Jaya (Fig. 7c, f, i). The changes in unsuberized cells on drought side of the root was more in Jaya and IR-20 than Pokkali and ARB6 (Fig. 7c, f, i), suggesting the increase of passage cells in tolerant varieties like ARB6 and Pokkali on the expense of suberized cells.

The change in the pattern of suberization followed the same trend in the exodermis, but less. The percentage of fully suberized cells in the exodermis appeared comparable across all varieties in both water and drought side of the root except Pokkali where it was more (Supplementary Fig. 1a, d, g). There was a significant increase in passage cells in mid and base region in both sides of the root of ARB6 and Pokkali with a concomitant decrease in the percentage of unsuberized cells (Supplementary Fig. 1b, e, h). The changes were more dramatic in ARB6. In all three regions analyzed – base, middle (mid) and tip of the root, IR-20 had a larger proportion of unsuberized cells in base and mid than the other three varieties, in tip region it was comparable across all four varieties (Supplementary Fig. 1c, f, i).

### Suberin deposition under W/S condition

The extent of suberization increased in salt stressed side of the root in all four varieties and in all regions of endodermis (Fig. 8a, d, g). An increase in passage cells in salt side of the root was seen in all varieties except Pokkali where there was a decrease (Fig. 8b, e, h). In base region, no unsuberized cells were seen in the endodermis of any variety except IR-20 (Fig. 8c, f, i), whereas unsuberized cells were observed in both mid and tip regions in all four varieties (Fig. 8c, f, i). The extent of suberization in exodermis was very less but followed the same trend as that of endodermis where suberized cells in Pokkali, ARB6 and Jaya developed on the expense of unsuberized cells (Supplementary Fig. 2).

**Figure 8.**
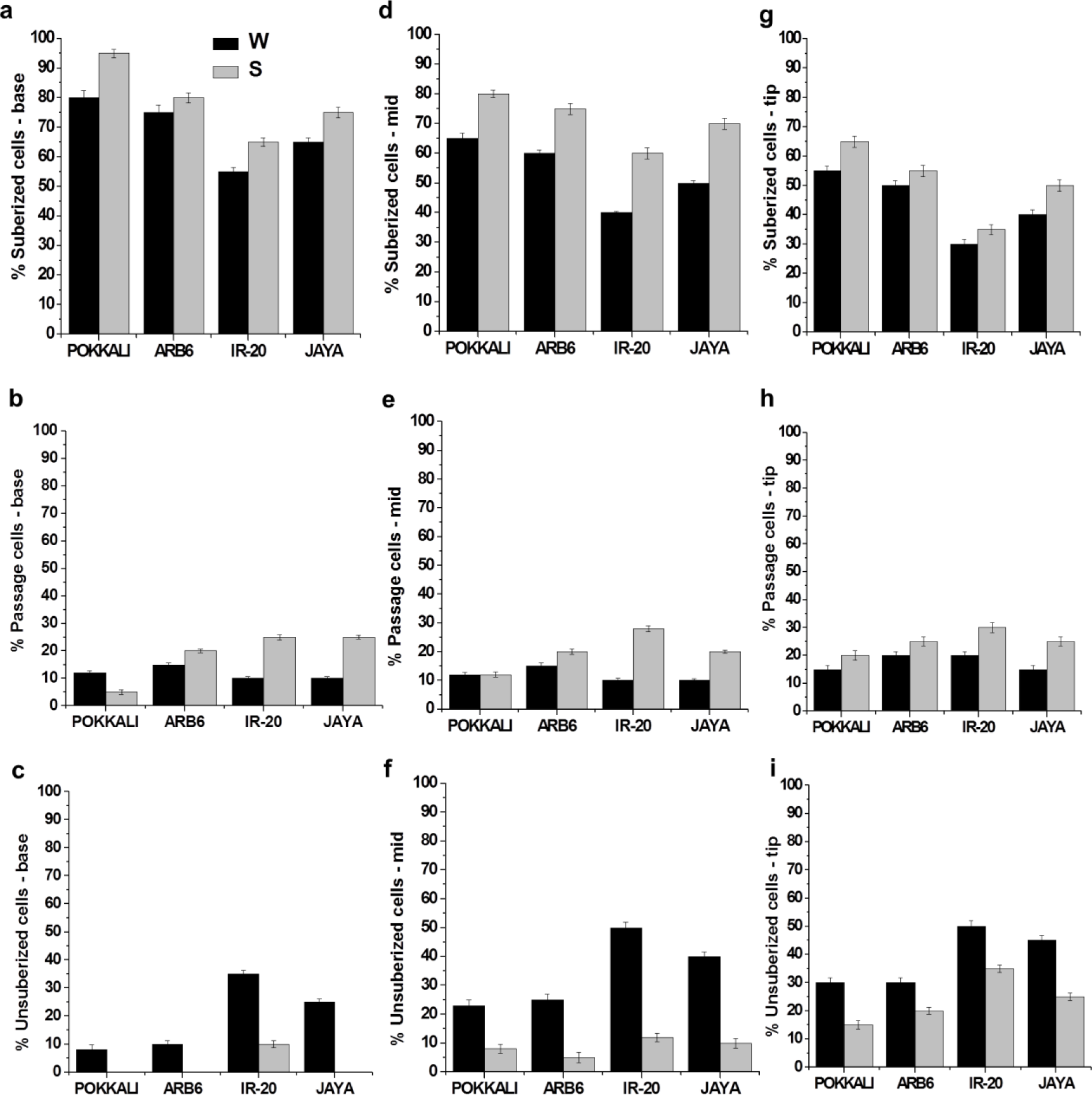
Quantification of suberization patterns under water/salt (W/S) condition at (a - c) base, (d - f) mid and (g - i) tip zone of the root endodermis to show suberized cells (having suberization on both sides and tangential areas), passage cells (with no tangential suberization) and unsuberized cells (without any suberin deposition). Data represents mean (±SE; n = 6)

### ABA in xylem sap

Roots synthesize ABA and transport it to shoots via xylem stream as an early signal or warning on experiencing abiotic stress like drought or salinity to prepare the plant for these stresses. To check how much ABA would be produced by the roots of four rice varieties when either sides or half of the root would be exposed to either drought or salinity or both. The ABA concentration in xylem sap under W/W condition from Day 1 to Day 7 was very low and comparable in all four varieties (Fig. 9a). Interestingly, the ABA concentration increased dramatically after Day 1 of exposure of these varieties to a combination of drought, salinity and well-watered condition (Fig. 9b, c, d, e, f). Highest concentration of ABA after Day 1 of stress was observed under D/S condition, followed by S/S, D/D, W/D and W/S condition in all four varieties (Fig. 9b, c, d, e, f), where the ABA concentration was always higher in ARB6 and Pokkali than the other two varieties. These results indicate that the roots under D/S, S/S and D/D condition, where both halves of these roots were either exposed to Drought or salinity or both, were exporting significant amount of ABA to shoots through xylem stream compared to W/D and W/S condition where only half of the root was either exposed to drought or salinity and the other half was in well-watered condition. It was also observed that the concentration of ABA in xylem sap collected after Day 2, Day 3, Day 4 and Day 7 of stress, declined with each day untill the end of the stress period, indicating that the roots of these varieties had presumably reduced the export of ABA to shoots compared to Day 1 under all conditions.

**Figure 9.**
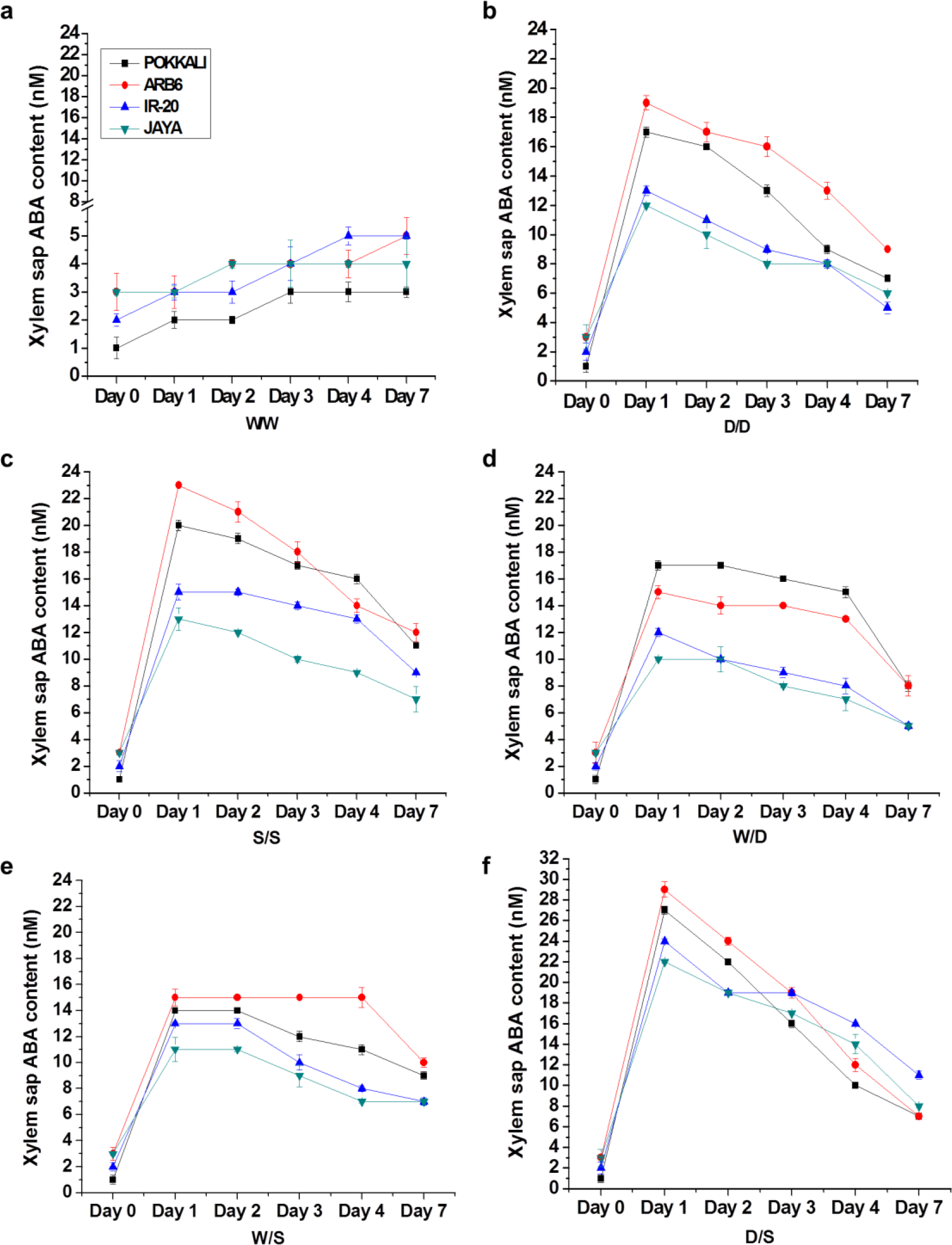
Abscisic acid **(ABA)** content in xylem sap of four rice varieties subjected to (a) water/water (b) drought/drought (c) salt salt (d) water/drought (e) water/salt and (f) drought/salt condition for a week in a split root experiment. Data represents mean ± SE, n = 3.

### ABA in leaves

Like xylem sap, the ABA concentration in leaves under W/W condition was also very low and comparable in all four varieties from Day 1 to Day 7 (Fig. 10a). The dramatic increase in concentration of ABA in leaves after Day 1 correlated with that of Xylem sap ABA under D/S, S/S, D/D, W/D and W/S conditions in all varieties, with ARB6 and Pokkali having high concentrations than Jaya and IR-20 (Fig. 10b, c, d, e, f). Interestingly, the ABA concentration estimated in leaves of Pokkali and ARB6 from Day 2 to Day 7 followed the same trend of decline as that of xylem sap ABA, indicating a direct connection between the ABA accumulated in leaves with that sent from roots (Fig. 10b, c, d, e, f). But surprisingly, such a trend was not observed in case of Jaya and IR-20, in fact, as the soil dried or became more saline, the ABA concentration from Day 2 to Day 7 in these two varieties was found to increase irrespective of the decline in xylem sap ABA (Fig. 10b, c, d, e, f). This would presumably indicate the shoot borne ABA accumulation in leaves of these two varieties.

**Figure 10.**
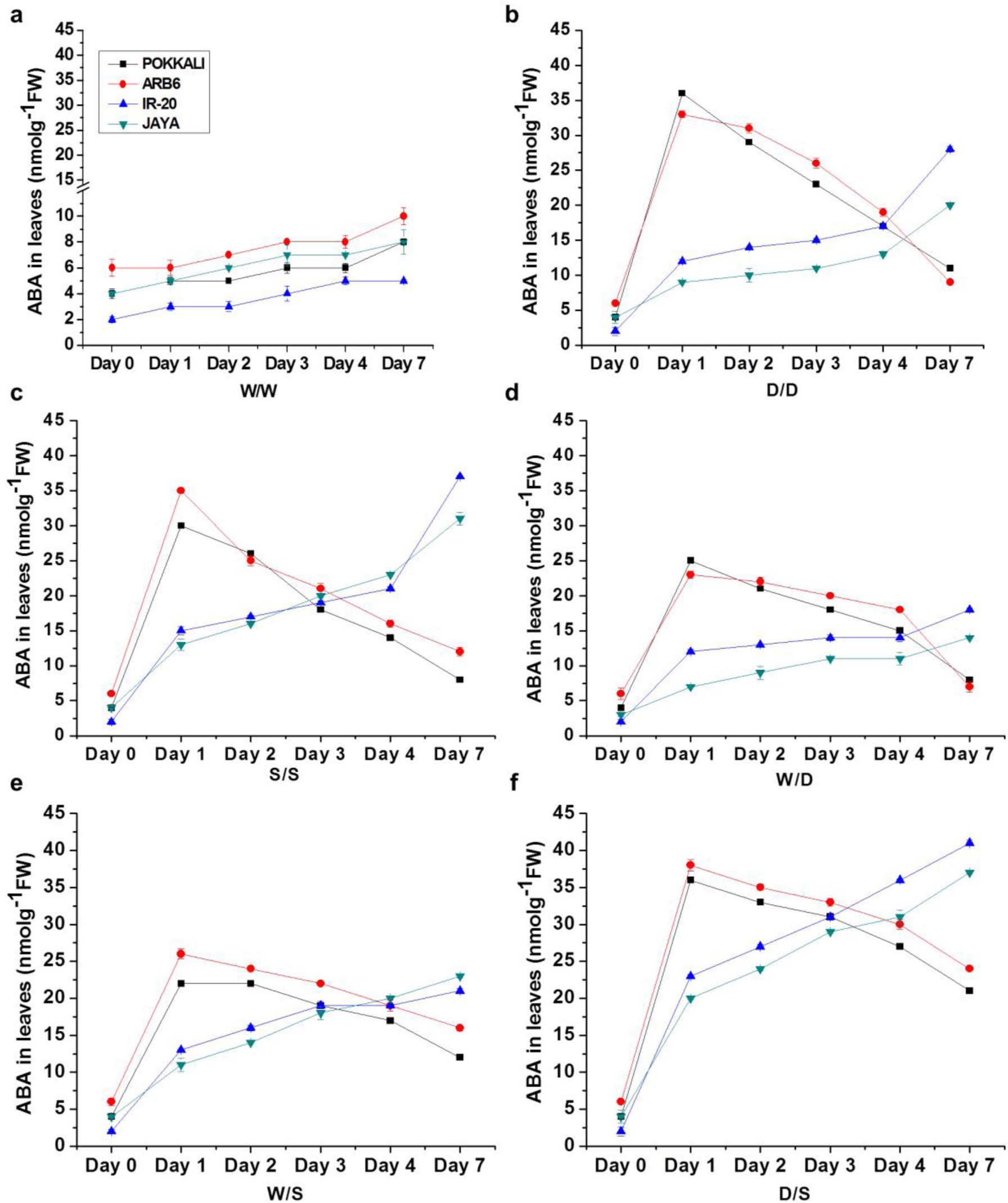
ABA content in leaves under (a) water/water (b) drought/drought (c) salt/salt (d) water/drought (e) water/salt and (f) drought/salt condition for a week. Data represents mean ± SE, n = 3.

### Stomatal conductance

There was no difference in the stomatal conductance from Day 0 to Day 7 under W/W condition in all four varieties, with ARB6 having high stomatal conductance than Pokkali, Jaya and IR-20 (Fig. 11a). Interestingly, the stomatal conductance in all four varieties under D/D, S/S, W/D, W/D and D/S conditions was not found in correlation with xylem sap ABA concentration (Fig. 11b, c, d, e, f), suggesting the stomatal conductance is independent of root produced ABA. Unlike tolerant varieties like Pokkali and ARB6, the stomatal conductance of Jaya and IR-20 was observed to be in correlation with ABA in leaves (Fig. 11b, c, d, e, f), suggesting that in tolerant varieties the stomatal conductance could be dependent on other factors like root length and osmotic adjustment (ability of roots to allow water movement).

**Figure 11.**
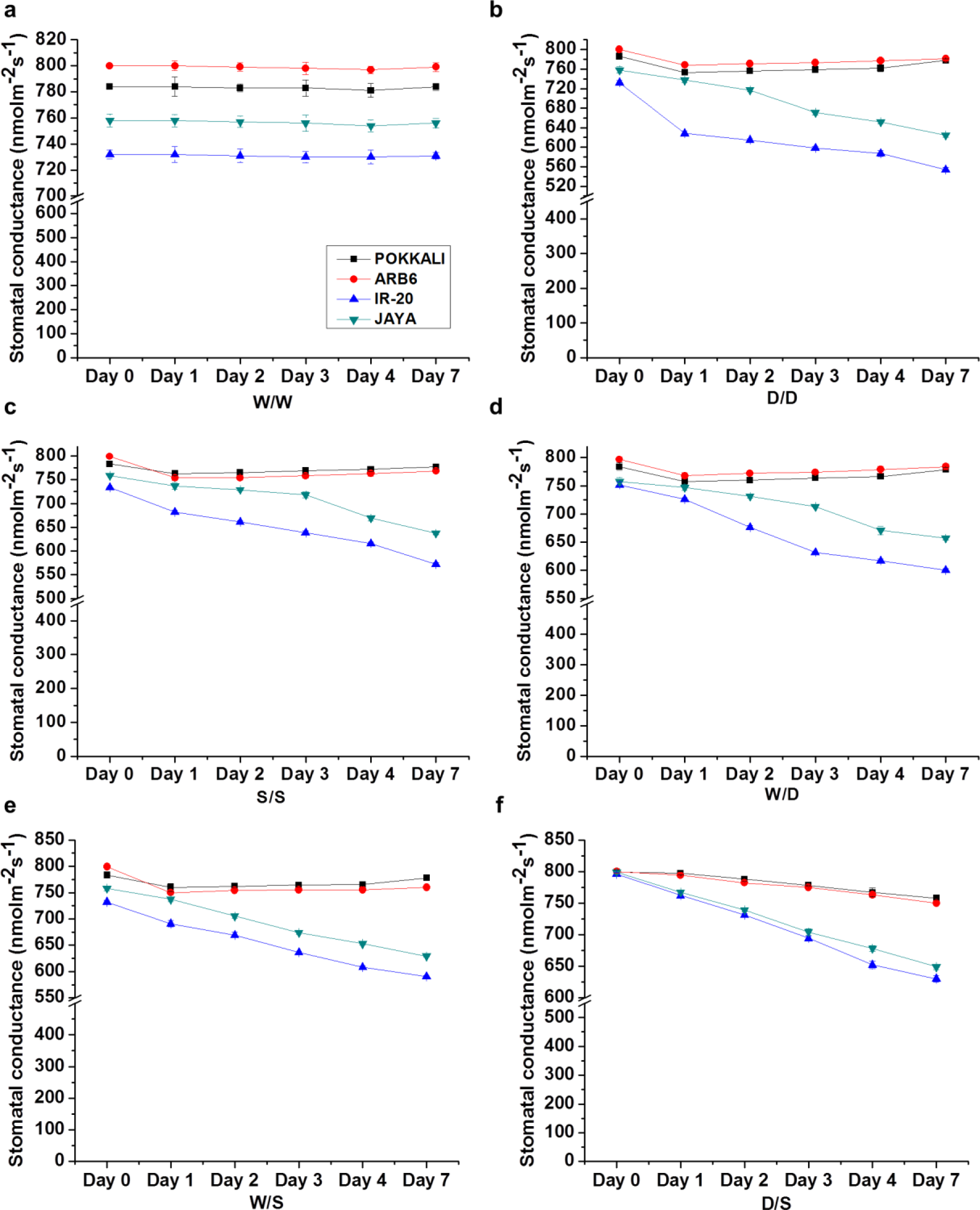
Stomata! conductance in four rice varieties subjected to (a) water/water (b) drought/drought (c) salt/salt (d) water/drought (e) water/salt and (f) drought/salt condition for a week. Data represents mean ± SE, n = 6.

## Discussion

Drought and salinity are considered two highly challenging stresses for rice plant. Role of roots is essential not only in perceiving the water deficit condition due to drought and salinity stress, but also in communicating this information to the shoot, which in turn helps plants to develop strategies to cope with water deficit condition either by mining water from deeper soil layers or by reducing the transpiration rates. Although both drought and salinity essentially impose osmotic stress but salinity has the added feature of ion toxicity (Munns and Tester, 2008; Gorham et al., 2009), therefore, to survive both stresses, root architecture and physiology may be expected to be different, as root traits necessary to cope with drought may not be promising to survival under salinity and vice versa. Although, the split-root system has been used to study plant response to heterogeneous soil conditions such as heterogeneous nutrient distribution (Arredondo and Johnson, 1999; Paterson et al., 2006), partial root drying (Lawlor, 1973; Sobeih et al., 2004) and unequal salt distribution (Shani et al., 1993; Messedi et al., 2004; Lycoskoufis et al., 2005; Bazihizina et al., 2009), this study was to test the effect of uniform and non-uniform drought and salinity stress on rice roots, and to understand as to how these roots would communicate and help tolerant and sensitive cultivars to cope with these stresses.

The finding that tolerant varieties like Pokkali and ARB6 had grown longer roots in drought and salt side of the PVC pipe under D/D, S/S, W/D, W/S and D/S condition (Fig. 3a) suggests that these roots on either side of the pipe are not only growing independent of each other but also in response to the stress imposed. Further it also suggests that these two varieties are resorting to water mining strategy by growing longer roots to overcome the water deficit condition under these stresses. However, the sensitive varieties Jaya and IR-20 preferred to grow roots well on water side of the pipe with no efforts on root growth on the stress side of the pipe. The observation of a large differential between canopy and ambient temperature in both ARB6 and Pokkali under W/W, D/D, S/S, W/D and W/S condition strongly suggests evapotranspirational cooling of canopy and a considerable transpirational stream even under D/S condition (Fig. 4a). High rates of xylem fluid exudation by both varieties under W/W, D/D, S/S, W/D and W/S condition (Fig. 4b) are consistent with observations of canopy temperature. Both Pokkali and ARB6 have long roots that elongate further under stress conditions, therefore, facilitating access to subsurface water which in turn helps maintaining normal transpiration in the face of water deficit condition. The water use of the whole plant was also reported to be higher when roots were exposed to non-uniform than high uniform salinity, and more water was found to be absorbed from the low salinity side (Shani et al., 1993; Bazihizina et al., 2009).

Under water deficit conditions, osmotic stress and high concentrations of salts in the soil make it difficult for roots to absorb water and therefore, can only grow if they osmotically adjust. Both in halophytes (Flowers and Colmer, 2008; Hariadi et al., 2011) and in glycophytes (Shabala and Lew, 2002), the increased uptake of inorganic ions is the most important means of osmotic adjustment. Indeed, both in ARB6 and Pokkali sap uptake under D/D, S/S, W/D, W/S and D/S condition was driven by accumulation of osmolytes (Fig. 5a, b, c, d, e). In ARB6, accumulation of osmolytes was much higher in the face of drought stress than in salinity while the opposite was observed for Pokkali. Jaya also accumulated a considerable amount of osmolytes under saline side of the root than under drought, which is consistent with the findings where Jaya was shown to adapt well to salinity (Krishnamurthy et al., 2009). In addition, the dwindling supply of water to one portion of the root system can be compensated by the other portion as shown in the study where well-watered root systems of apple (West, 1978) or tomato (Papadopoulos et al., 1985) compensated for the dwindling supply of water by salinized roots. The roots on water side of both W/D and W/S could very well be compensating for the inadequate supplies of water on the other half, which is why the sap exudation rates under D/S condition were observed to reduce to a great degree.

Besides osmolyte accumulation, pattern of suberization also attributes to the root hydraulics to a great extent (Clarkson and Robards, 1975; Melchior and Steudle, 1993). Indeed, the impressive sap uptake of Pokkali and ARB6 under W/D condition must be ascribed to decreased suberization and increased passage cell number in both endodermis and exodermis on drought side of the root (Fig. 8 and Supplementary Fig. 1). Also, under W/S condition, enhanced suberized cell number in both endodermis and exodermis on the expense of unsuberized cells was observed in salt side of the root in Pokkali than the other three varieties (Fig. 9 and Supplementary Fig. 2), while the plants were still able to maintain the considerable number of passage cells. Presence of passage cells ensures enhanced fluid flow through the symplastic pathway under both drought and salinity condition, while excellently avoiding the uptake of toxic Na^+^ ions in latter. These findings of suberization patterns in salinity and drought of these four varieties very well correspond to the observations of (Krishnamurthy et al., 2009; Chowdery et al., unpublished data) in non-split root studies.

The balance of ABA biosynthesis and catabolism controls the plant endogenous ABA levels (Zeevaart and Creelman, 1988). When the water potential of the leaves is affected, ABA accumulates in root tissue, then is released into the xylem and finally transported to the shoot (Zhang and Davies, 1990; Davies et al., 2005), which in turn regulates the stomatal conductance in leaves (Davies and Zhang, 1991). Having acted on the stomata ABA is rapidly metabolized and not accumulated in the leaf (Peuke et al., 1994; Peuke et al., 2002).

ABA concentration estimated in the xylem sap collected during 7 days of exposure to uniform and non-uniform water, drought and salinity condition, was observed to increase dramatically after Day 1 in all four varieties, with more increase in Pokkali and ARB6 than the other two varieties under all conditions except W/W condition (Fig. 10). Interestingly, more increase in xylem sap ABA was found under D/S, S/S and D/D condition than W/D, W/S and W/W condition. The xylem sap ABA levels decreased with each day until the end of the stress period. Interestingly, the ABA levels in leaves of tolerant varieties like Pokkali and ARB6 followed the same trend of initial increase followed by decrease as that of xylem sap ABA, suggesting the control of latter on levels of ABA in leaves in these two varieties. However, the opposite trend was shown by Jaya and IR-20, despite the decreasing levels of xylem sap ABA, the ABA concentration in leaves kept increasing, suggesting a possible contribution of leaf borne ABA accumulation in the leaves in these two varieties. During drought and salinity condition, root cells experience the osmotic stress due to lower water status which in turn could activate mechano-sensitive Ca^2+^ channels and have profound effects on ion transport systems such as K^+^-H^+^ symport (Netting, 2000) and eventually resulting in the changes in the pH of xylem sap. It is possible that an increase in pH could have contributed to the increase of ABA within leaves of Jaya and IR-20. An increase in xylem pH was shown to reduce the catabolism of ABA and therefore cause an increase in ABA synthesis in leaves (Wilkinson and Davies, 1997). The stomatal conductance of Pokkali and ARB6 showed no change while that of Jaya and IR-20 corresponded well with the increased levels of Leaf ABA, suggesting that leaf borne ABA indeed controls the stomatal conductance in sensitive varieties and is independent of the root sourced ABA. The results are in agreement with the findings where using reciprocal grafting between wild-type and mutant plants, the stomatal conductance during water stress was reported to be determined by the shoot rather than the root genotype in tomato (Holbrook et al., 2002) and sunflower (Fambrini et al., 1995).

Since Jaya and IR-20 do not possess long root system to fetch water from deeper layers of drying or saline soil as in case of D/D, S/S, W/D, W/S and D/S condition, they resort to the strategy of water conservation by leaf borne ABA induced stomatal closure (Lee and Luan, 2012; Bauer et al., 2013). Closing the stomata under water deficit conditions assures water status of the plant, but the amount of water lost through the leaves in tolerant Pokkali and ARB6 is well balanced by the amount taken up by the roots because they possess longer roots with numerous passage cells and an added feature of better osmotic adjustment which ensures adequate water mining from subsurface soils.

## Conclusion

Our split root study with rice under uniform and non-uniform water, drought and salinity conditions provided excellent understanding about the growth and physiological response of rice roots to heterogeneous soil conditions. Both halves of the root on either side of the PVC pipe growing in different conditions like W/D, W/S and D/S, responded independent of the other half in terms of osmotlyte accumulation and pattern of suberization. It was observed that in rice ABA signals are produced from roots in response to drying and saline soil and transported through xylem sap. The stomatal conductance is not related to xylem sap ABA concentration. Also, the data presumes either the signal from root, pH change or in-situ synthesis caused leaf ABA level to increase which inturn controls the stomatal conductance.

## Acknowledgements

We are grateful to Department of Plant Biotechnology, University of Agricultural Sciences, Bangalore to provide the field station and timely resources to conduct these experiments. The work was performed with internal NCBS funding. The Central Imaging & Flow Cytometry Facility (CIFF) at NCBS Bangalore is gratefully acknowledged.

