## Supplemental Figures for "Highly Localized Responses to Salinity and Drought in Rice Roots"

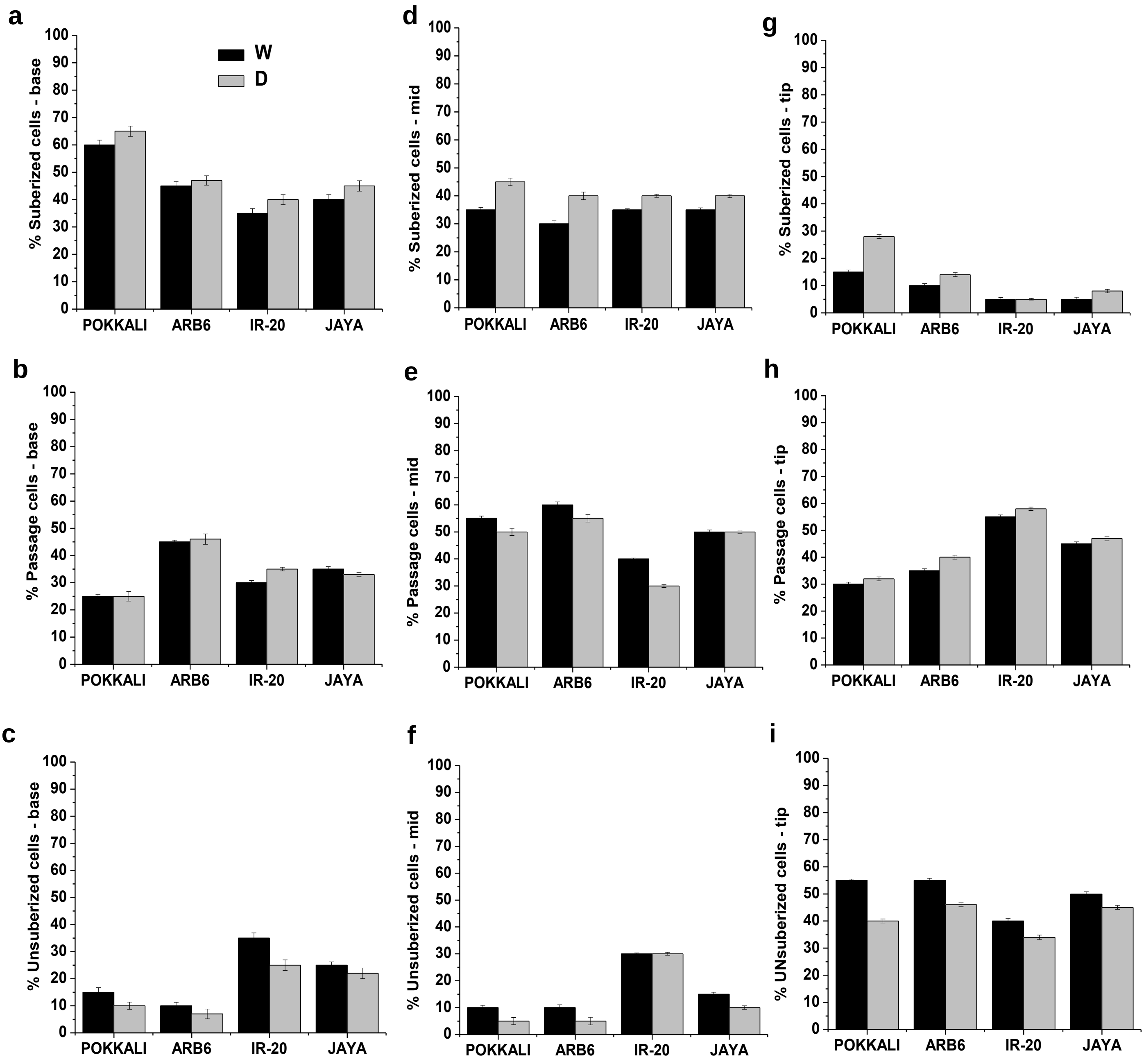


**Supplementary Figure 1** Quantification of suberization patterns in (a - c) base, (d - f) mid and (g - i) tip zone of the root exodermis to show suberized cells (having suberization on both sides and tangential areas), passage cells (with no tangential suberization) and unsuberized cells (without any suberin deposition) under water/drought (W/D) condition. Data represents mean (±SE; n = 6)

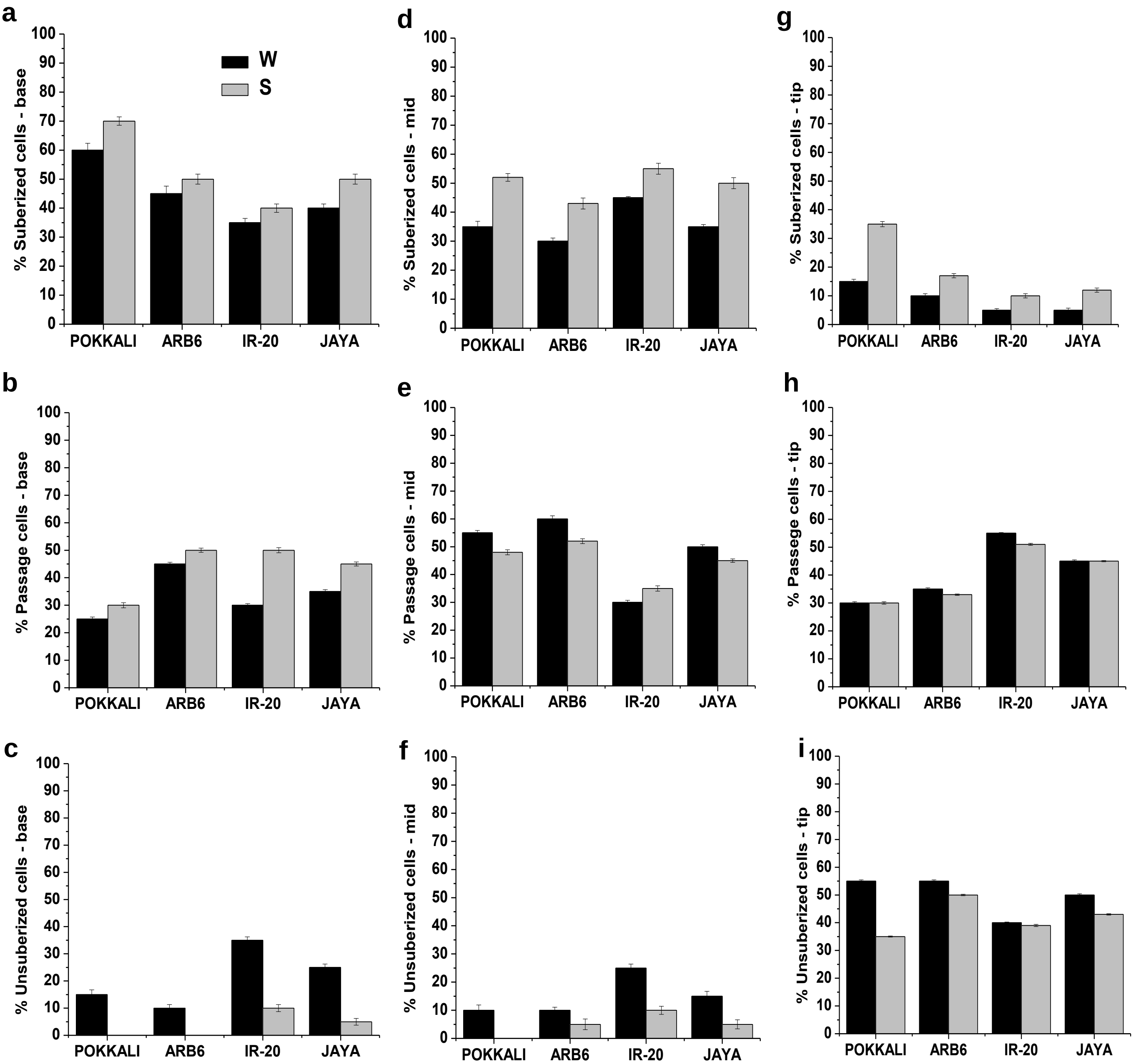


**Supplementary Figure 2** Quantification of suberization patterns in (a - c) base, (d - f) mid and (g - i) tip zone of the root exodermis to show suberized cells (having suberization on both sides and tangential areas), passage cells (with no tangential suberization) and unsuberized cells (without any suberin deposition) under water/salt (W/S) condition. Data represents mean (±SE; n = 6)
